# Homeostatic mechanisms regulate distinct aspects of cortical circuit dynamics

**DOI:** 10.1101/790410

**Authors:** Yue Wu, Keith B. Hengen, Gina G. Turrigiano, Julijana Gjorgjieva

## Abstract

Homeostasis is indispensable to counteract the destabilizing effects of Hebbian plasticity. Although it is commonly assumed that homeostasis modulates synaptic strength, membrane excitability and firing rates, its role at the neural circuit and network level is unknown. Here, we identify changes in higher-order network properties of freely behaving rodents during prolonged visual deprivation. Strikingly, our data reveal that pairwise functional correlations and their structure are subject to homeostatic regulation. Using a computational model, we demonstrate that the interplay of different plasticity and homeostatic mechanisms can capture the initial drop and delayed recovery of firing rates and correlations observed experimentally. Moreover, our model indicates that synaptic scaling is crucial for the recovery of correlations and network structure, while intrinsic plasticity is essential for the rebound of firing rates, suggesting that synaptic scaling and intrinsic plasticity can serve distinct functions in homeostatically regulating network dynamics.

## Introduction

Neural circuits are faced with a fundamental problem: how to allow experience to alter and refine network connectivity during learning and experience-dependent plasticity, while still maintaining stability of function. Generating a neural system that is both stable and flexible is a non-trivial challenge and requires a prolonged period of development when multiple mechanisms at the level of single neurons and networks of neurons interact. Two powerful and fundamentally different forms of plasticity involved in this process are Hebbian mechanisms, which alter synaptic connectivity in a synapse-specific manner, and homeostatic mechanisms that maintain stable function by globally adjusting overall synaptic weights and neuronal excitability.

The development and refinement of visual response properties in primary visual cortex (V1) involves classic synapse-specific mechanisms implementing the bidirectional form of Hebbian plasticity, such as long-term potentiation (LTP) and long-term depression (LTD), considered to be the cellular substrate for learning and memory (Smith et al., 2008). However, associative Hebbian plasticity drives positive feedback processes leading to unstable network dynamics and some form of homeostasis is needed to compensate for the inherent instability (Abbott and Nelson, 2000; Turrigiano and Nelson, 2004). A large body of evidence shows that various homeostatic plasticity mechanisms, including synaptic scaling and intrinsic plasticity (Turrigiano et al., 1998; Desai et al., 1999), operate in the brain to maintain stability despite various internal and external perturbations. More specifically, homeostatic plasticity can elevate neural activity in response to sensory deprivation (Hengen et al., 2013; Keck et al., 2013), and suppress activity under conditions of overexcitation (Seeburg et al., 2008; Evers et al., 2010).

Despite great efforts to describe homeostatic mechanisms at the single cell level, how network properties are homeostatically regulated is largely unknown. Furthermore, while Hebbian and homeostatic mechanisms operate at different timescales and can be induced by distinct cues (Stellwagen and Malenka, 2006; Shepherd et al., 2006; Joseph and Turrigiano, 2017; Daoudal and Debanne, 2003), how they interact within complex, highly recurrent microcircuits, as those found in the cortex, to refine and maintain circuit function has remained elusive. A critical challenge has been the lack of detailed measurements of individual synaptic strengths and their potential impact on large-scale network dynamics, especially in a highly recurrent network like the cortex.

Here we investigate two main questions. First, which aspects of network function are under homeostatic control? Second, why are there so many homeostatic mechanisms, and do they serve redundant or unique functions? To address these questions, we combine analysis of *in vivo* electrophysiological data during sensory deprivation in rodent visual cortex and computational modeling of cortical synaptic plasticity and network dynamics. First, we analyzed the collective activity of multiple neurons in the monocular region of primary visual cortex (V1m) during a classic monocular deprivation (MD) paradigm (lid suture), in freely behaving rats over 9 days during the critical period (Hengen et al., 2016). Earlier work demonstrated that MD induces an initial drop in firing followed by the rates’ homeostatic recovery despite long-lasting deprivation (Hengen et al., 2016). Here we reanalyzed these datasets to characterize the temporal evolution of higher-order network properties over the same nine-day period. Individual pairwise correlations, including correlation structure, weakened during brief MD, but recovered during prolonged MD. Second, to understand how the cortical network exploits diverse homeostatic mechanisms to return firing rates and correlations to baseline after prolonged MD, we took advantage of a plastic spiking recurrent network model equipped with known plasticity and homeostatic mechanisms. Our work suggests that synaptic scaling is crucial for the recovery of correlations and network structure, whereas intrinsic plasticity is essential for the rebound of firing rates. These results indicate that different homeostatic mechanisms act in the brain to independently regulate distinct network features.

## Results

### Pairwise correlations during the critical period and in response to monocular deprivation

We first confirmed previous analysis of individual neurons recorded *in vivo* in the primary visual cortex during the critical period of plasticity (postnatal days 24 to 32). In these experiments, MD was performed after 3 days of baseline activity and continued for the rest of the recordings. While firing rates of individual neurons remain relatively stable during normal development, brief two-day monocular deprivation (MD) caused the firing rates to decrease to 40% of their baseline values (Fig. 1A, left) (Hengen et al., 2013, 2016). However, despite prolonged MD, over the next three to four days firing rates gradually recovered to baseline after an initial overshoot (Fig. 1A, right) (Hengen et al., 2013, 2016). These effects were not only observed at the population level, but also at the level of individual neurons (Hengen et al., 2016). Here, we investigated higher-order network properties during normal development and following prolonged MD by calculating the next statistical moment beyond the firing rates, namely the pairwise spiking correlations between different neuron types (Methods). Specifically, we quantified the temporal evolution of the correlation coefficient of individual neuron pairs and of the average correlation across all pairs both during normal development and after perturbing visual input through MD. In control hemispheres, unlike firing rates, correlations increased slightly as a function of age (n = 5 animals, Fig. 1B, left). By contrast, in deprived hemispheres, correlations initially dropped over the first two days and then gradually rebounded to pre-deprivation levels (n = 5 animals, Fig. 1B, right), displaying a similar pattern as the firing rates (Fig. 1A). As previously reported, we observed light-dark oscillations in the correlation amplitudes with higher correlations in the light and lower correlations in the dark (Pacheco et al., 2019).

**Figure 1:**
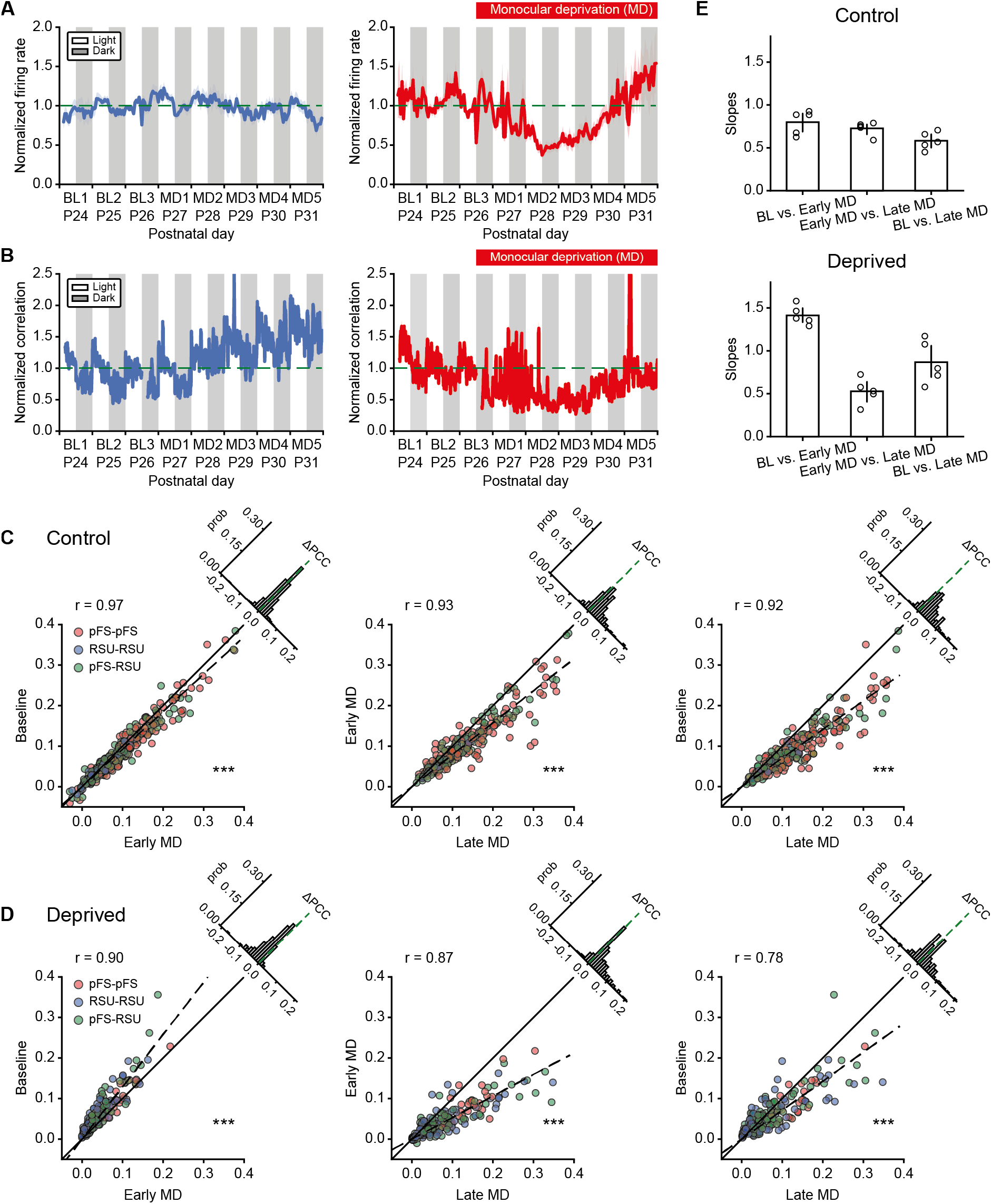
MD induces an initial drop in correlations followed by their homeostatic recovery. **A**. The average firing rates of 80 neurons from five control hemispheres (left) and 104 neurons from five deprived hemispheres (right) normalized to the firing rates at P26 in the light (horizontal dashed line). **B**. The average pairwise correlations of 970 pairs from five control hemispheres (left) and 2455 pairs from five deprived hemispheres (right) normalized to the correlations at P26 in the light (horizontal dashed line). **C**. Correlation comparisons between baseline (BL) and early MD (left), between early MD and late MD (middle), and between BL and late MD (right) at the single cell-pair level of one control hemisphere. Different colors represent the correlations between different neuron types. Dashed lines are fitted regression lines crossing the origin. Upper left histograms indicate the distributions of correlation differences. *** p *<* 0.001, Wilcoxon signed-rank test. **D**. Same as C but for one deprived animal. Here, for two hemispheres we used MD3 and for the other three hemispheres MD2 as early MD because different animals showed the biggest drop in correlations at different times. *** p *<* 0.001, Wilcoxon signed-rank test. **E**. Slopes of fitted regression lines for the correlation comparisons in C and D. Data are shown as means *±* SEM.

To assess the degree to which correlations of individual neuron pairs changed beyond the population level, we evaluated single cell-pair correlations on different days. As in earlier analysis, neurons were separated into putative PV+ fast-spiking units (pFS) or regular-spiking units (RSUs) based on waveform and spiking characteristics (Hengen et al., 2013, 2016). Specifically, we focused on three different 12-hour periods recorded in the light: (1) baseline (BL) corresponding to P26, (2) a period that we called “early MD” when the largest drop of firing rates and correlations occurred, typically two or three days after baseline i.e. P28 or P29, and (3) a period that we called “late MD” corresponding to the time when the firing rates and correlations nearly recovered to baseline i.e. P31. As observed already for the average correlation combining all neuron pairs and animals, correlations increased during normal development covering the 6-day period during which recordings were performed. The increase between BL and early MD was small (n = 435 pairs, Fig. 1C, left, *r* = 0.97, *p <* 10^*−*21^, Wilcoxon signed-rank test). Correlations at late MD were significantly greater than at early MD (n = 253 pairs, Fig. 1C, middle, *r* = 0.93, *p <* 10^*−*28^, Wilcoxon signed-rank test). The developmental increase in correlations during the critical period became most obvious when we compared BL versus late MD (n = 253 pairs, Fig. 1C, right, *r* = 0.92, *p <* 10^*−*37^, Wilcoxon signed-rank test). We did not observe any obvious differences in correlations among different cell types in that they all showed similar patterns of temporal evolution. Moreover, almost all neuronal pairs in the control hemispheres demonstrated an increase in correlation (Fig. 1C, right).

Conversely, in deprived hemispheres, correlations of the majority of individual cell pairs, independent of their type, underwent an initially significant drop during early MD (n = 190 pairs, Fig. 1D, left, *r* = 0.90, *p <* 10^*−*24^, Wilcoxon signed-rank test) followed by a subsequent increase during late MD (n = 231 pairs, Fig. 1D, middle, *r* = 0.87, *p <* 10^*−*27^, Wilcoxon signed-rank test). The correlations during late MD recovered to a higher level than baseline (n = 190 pairs, Fig. 1D, right, *r* = 0.78, *p <* 10^*−*4^, Wilcoxon signed-rank test). We summarized the gradual increase of correlations in control hemispheres and drop followed by recovery in deprived hemispheres by the fitted regression lines of the individual pair data for each animal (Fig. 1E). Remarkably, despite a degree of variability across animals, the drop and recovery of correlations induced by MD were ubiquitous (Fig. S1).

### Network structure after monocular deprivation

While correlations at the single cell-pair level recovered during late MD, the difference between correlations at late MD and BL (Fig. 1D, right) raised the possibility that the recovered network might have a different structure after recovery. To examine the evolution of network structure during normal development over the critical period and during prolonged MD, we examined the correlation matrices on different days. An example experiment shows that in the control hemisphere, the structure of the correlation matrix remained consistent over time (n = 11 neurons, Fig. 2A), whereas in the deprived hemisphere, the correlation structure initially weakened and recovered to a similar structure as baseline (n = 14 neurons, Fig. 2B). To quantify the similarity between the structure of correlation matrices at distinct time points, we used a metric known as the L1 distance (Methods), which measures the average positive distance between the vectors of correlations in a high dimensional space. This metric allowed us to summarize the results for multiple animals, and revealed that in control hemispheres the correlation matrix at BL is more similar to the correlation matrix at late MD relative to randomly shuffling the latter (n = 609 pairs, Fig. 2C, left, *p <* 10^*−*28^, Wilcoxon signed-rank test). Similarly, in deprived hemispheres, the correlation matrix at BL was more similar to the correlation matrix at late MD relative to randomly shuffling the latter (n = 505 pairs, Fig. 2C, right, *p <* 10^*−*30^, Wilcoxon signed-rank test). This demonstrates a high degree of similarity between correlation structure at baseline and at late MD. Interestingly, the correlation matrices were composed of several assemblies – groups of neurons exhibiting stronger pairwise correlations (Fig. 2A, B) – reminiscent of clustered network structure reported in previous studies (Ko et al., 2011; Cossell et al., 2015; Perin et al., 2011; Yoshimura et al., 2005).

**Figure 2:**
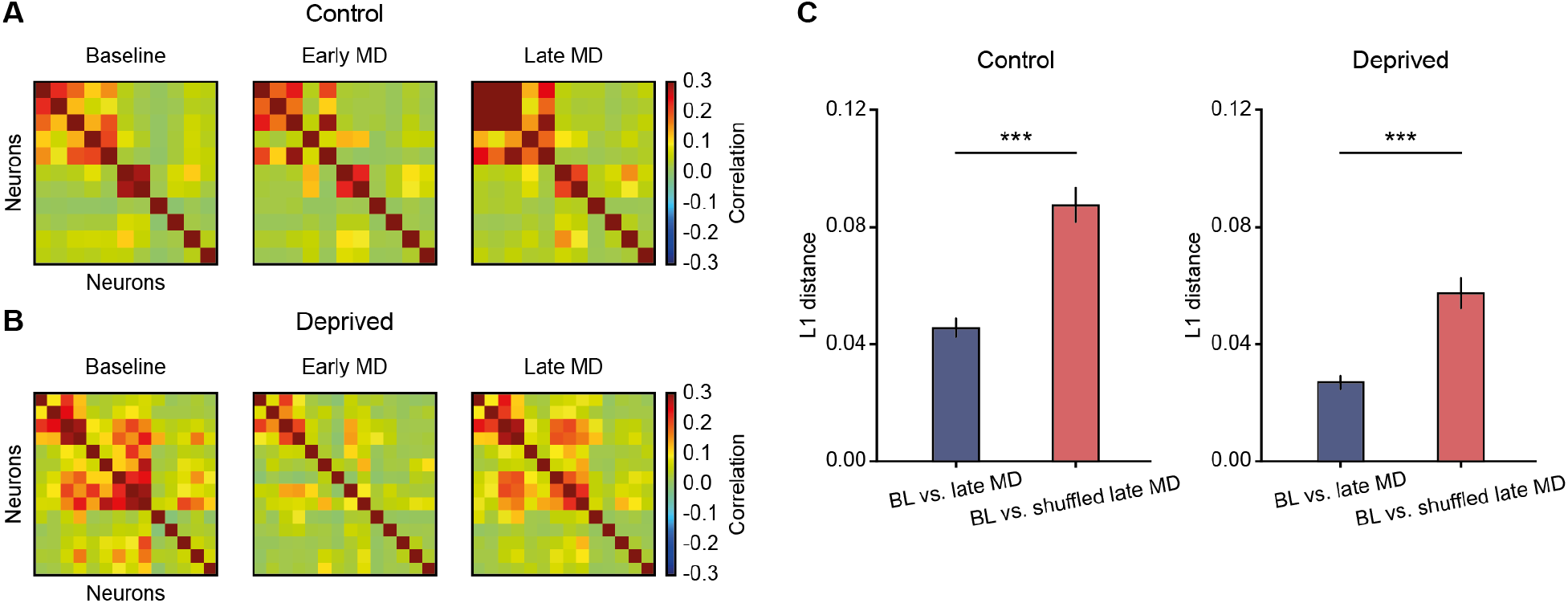
Structure of correlation matrices is maintained after recovery. **A**. Example correlation matrix of 11 neurons from one control hemisphere at three different time points. **B**. Same as A, but of 14 neurons from one deprived hemisphere. **C**. L1 distance between correlation matrix at baseline and at late MD, relative to baseline and shuffled late MD for control and deprived hemispheres. Data are shown as means *±* SEM. *** p *<* 0.001, Wilcoxon signed-rank test.

In conclusion, our analysis of V1 cortical activity recorded *in vivo* demonstrates that the correlation structure of these networks is homeostatically regulated following perturbation of normal sensory experience.

### Formation of structured connectivity assemblies during training in a recurrent network model

We next asked what mechanisms underlie the observed neuronal and network-level changes during normal development, and following a perturbation like MD. To understand how neural circuits exploit various synaptic plasticity and homeostatic mechanisms to decrease and recover both firing rates and correlations during MD, we built a plastic recurrent network model consisting of randomly connected excitatory and inhibitory spiking neurons (Methods). Model neurons receive thalamic inputs, with thalamocortical synaptic efficacy onto inhibitory neurons set higher than onto excitatory neurons, consistent with previous experimental studies (Cruikshank et al., 2007; Ji et al., 2015; Miska et al., 2018). Neuronal and network parameters were chosen to generate *in vivo*-like firing rates, with excitatory neurons firing at 5 Hz and inhibitory neurons firing at 13 Hz (Hengen et al., 2013).

To generate clustered correlation structure as observed experimentally (Fig. 2A, B), we followed previous modeling studies and included several experimentally characterized plasticity mechanisms (Litwin-Kumar and Doiron, 2014) (Methods). We first tasked the network with the imprinting of connectivity assemblies starting from an initially random connectivity. In contrast to previous computational modeling studies that used random, uncorrelated Poisson inputs (Litwin-Kumar and Doiron, 2014) and in line with our observation that the networks show stronger pairwise correlations in the light than in the dark (Pacheco et al., 2019), we postulated that input correlations – as would be generated during natural vision – matter for the generation of clustered connections. Therefore, we trained the network by stimulating the recurrent cortical network with thalamocortical Poisson spiking inputs that had identical firing rates, but differed in their correlation structure.

For the training, excitatory neurons were randomly grouped into 4 assemblies, although the exact number of assemblies was not important.

Before training with correlated inputs, the initial synaptic connections in the entire network were weak and identical between any pair of neurons of the same type (Fig. 3A, left), resulting in asynchronous irregular network activity (Fig. 3A, middle), and low correlations (Fig. 3A, right). During training, neurons within the targeted assembly received correlated inputs (Methods), which strengthened connectivity between them through recurrent Hebbian plasticity. After training, the network became structured with stronger synaptic connections within assemblies (Fig. 3B, left). As a result of this structure, the network no longer exhibited asynchronous irregular activity but blocks of activity defined as occasional periods of high firing rate (Fig. 3B, middle). The structured connectivity and block activity selectively increased correlations within assemblies (Fig. 3B, right).

**Figure 3:**
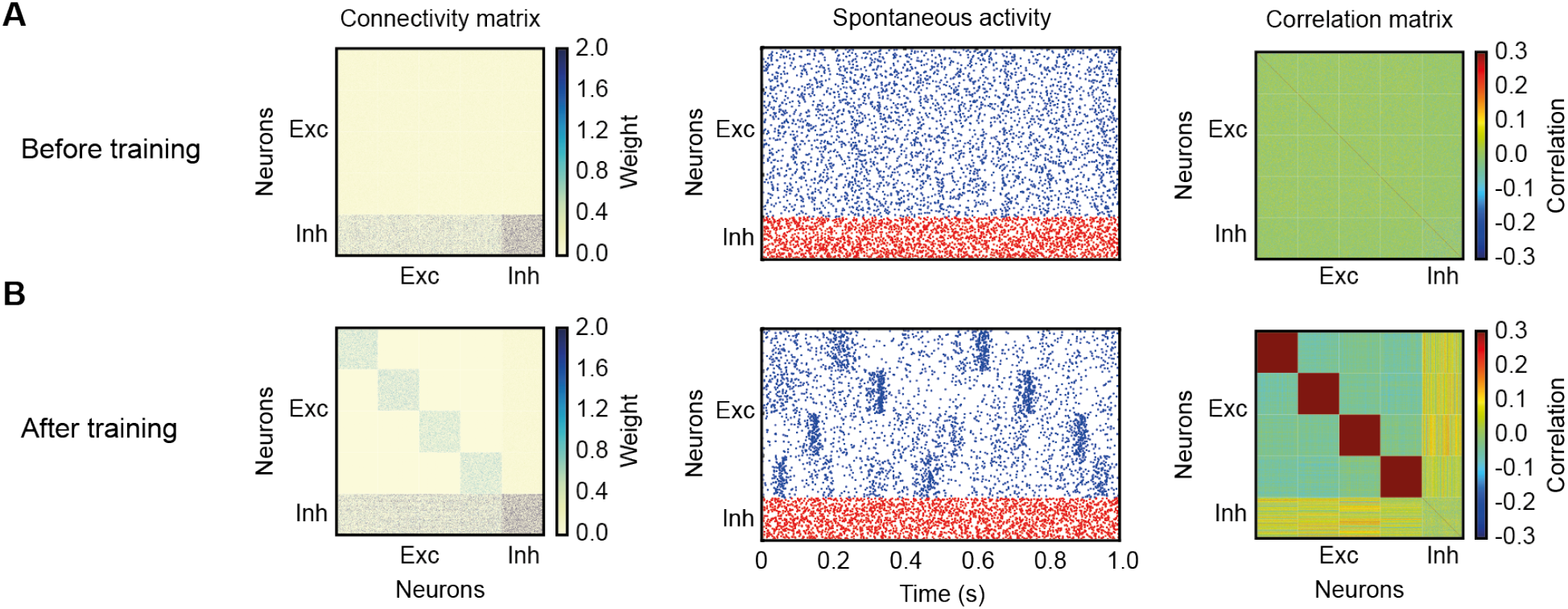
Imprinting connectivity assemblies with correlated inputs. **A**. Left: connectivity matrix before training. Middle: spontaneous activity of excitatory (blue) and inhibitory (red) neurons before training. Right: correlation matrix before training. **B**. Same as A, but after training.

### A model with persistent Hebbian LTD and homeostatic plasticity cannot recover correlations after MD

Using the structured model network as a baseline following normal cortical development after eye opening, we next wanted to investigate how this network responds to a sensory perturbation resembling MD. To achieve this, we needed to know how the inputs to the network are modified during MD. Previous experimental studies have reported that MD induces no change in the average firing rates of LGN, the visual area of the thalamus (Linden et al., 2009). Therefore, to simulate MD in our model network, we kept the firing rates of LGN inputs identical to that at baseline, but assumed that eye closure during MD considerably diminished input correlations. In the model, the excitatory neurons received uncorrelated Poisson inputs to denote the start of MD (Fig. 4A).

**Figure 4:**
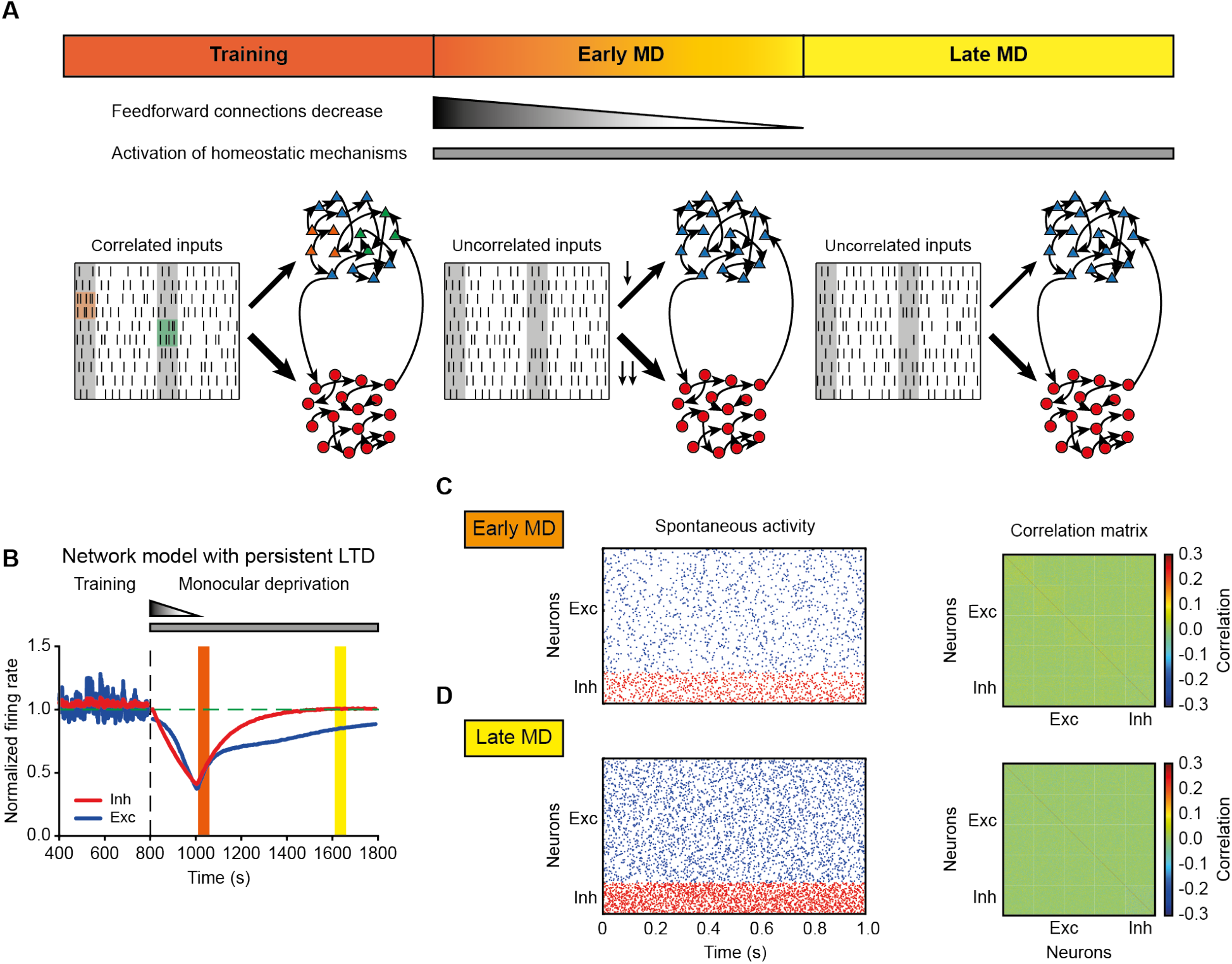
The model with persistent Hebbian LTD and homeostatic plasticity fails to recover correlations. **A**. Schematic description of the modeling framework during MD. The network consists of 80% excitatory neurons (triangles) and 20% inhibitory neurons (circles). Both of them receive thalamic inputs, and thalamocortical connections onto inhibitory neurons are stronger than onto excitatory neurons. Dashed lines indicate plastic connections, while solid lines represent fixed connections. Neurons receive correlated input at baseline (left), but uncorrelated input during early MD (middle) and late MD (right). During early MD, thalamocortical connections onto both excitatory and inhibitory neurons are linearly depressed, with thalamocortical connections onto inhibitory neurons depressed more strongly. The connections remain fixed during late MD. **B**. The average normalized firing rate of excitatory (blue) and inhibitory (red) neurons. The vertical dashed line indicates the onset of MD. The horizontal dashed line indicates a normalized firing rate of 1.0. **C**. Left: spontaneous activity of excitatory (blue) and inhibitory (red) neurons during early MD. Right: correlation matrix during early MD indicated by the orange region in B. **D**. Same as C but during late MD indicated by the yellow region in B.

In addition to these changes in input correlations, recent experiments have revealed that brief MD (2 days) induces long-term depression at thalamocortical synapses onto excitatory and inhibitory neurons, with thalamocortical synapses onto inhibitory neurons depressed more than synapses onto excitatory neurons (Miska et al., 2018). The process of long-term depression is not instantaneous, so we assumed that synaptic connections from thalamus to excitatory and inhibitory neurons undergo a linear decrease during the first two days of MD. To match experimental findings, the decrease in thalamocortical connections onto inhibitory neurons was larger (Fig. 4A, Methods). It is currently unknown when during MD this thalamocortical depression saturates, but since deprived-eye responsiveness reaches its minimum 2-3 days after the onset of MD (Frenkel and Bear, 2004), we assumed that the feedforward connections did not further decrease after this point, while keeping the inputs uncorrelated for the entire MD (Fig. 4A).

How does the recurrent network respond to these changes in input correlation structure and depression of feedforward connectivity strength that occur following MD? Although there are potentially multiple ways to achieve network stability and regulate network function, there are two fundamentally different mechanisms that are known to homeostatically control neuronal firing: homeostatic adjustment of synaptic strengths and of intrinsic excitability (Turrigiano and Nelson, 2004; Turrigiano, 2011; Lambo and Turrigiano, 2013). Neurons can regulate their activity by scaling incoming synaptic strengths in response to perturbations – a process known as synaptic scaling (Turrigiano et al., 1998). This scaling is bidirectional in that it can increase and decrease synaptic strengths, it is global and operates in a multiplicative manner. In addition to synaptic scaling, neurons can alter the number of ion channels to adjust their intrinsic excitability, and consequently modify their firing thresholds, in response to perturbations (Desai et al., 1999; Daoudal and Debanne, 2003; Grubb and Burrone, 2010).

Based on these experimental findings, besides Hebbian plasticity during training, we modeled these two distinct homeostatic mechanisms following MD: (1) synaptic scaling which acts only on excitatory synapses (Turrigiano et al., 1998; Hengen et al., 2013), and (2) intrinsic plasticity which modifies the intrinsic excitability of both excitatory and inhibitory neurons (Grubb and Burrone, 2010; Campanac et al., 2013) (Methods). In the presence of persistent thalamocortical LTD as during training, and both homeostatic mechanisms, the average firing rates of excitatory and inhibitory neurons in the model network first decreased to 40% of baseline, because slow homeostatic mechanisms could not overcome the feedforward synaptic depression and input decorrelation to recover firing rates. At the time that feedforward LTD saturated, firing rates started to increase due to homeostatic plasticity, resembling the recovery to baseline observed experimentally during late MD (compare Fig. 1B and Fig. 4B).

Next, we investigated the evolution of higher-order aspects of network dynamics. Similar to the analysis of our data, we focused on two key time points after MD onset in the model: early MD corresponding to the largest drop of firing rates (Fig. 4B, orange), and late MD corresponding to the time when the firing rates recovered close to baseline (Fig. 4B, yellow). The network showed irregular spiking dynamics with different firing rates during these two periods (Fig. 4C, D, left).

The correlations first decreased during the period modeling early MD as observed experimentally (compare Fig. 1B to Fig. 4C, right), but did not recover during the period corresponding to late MD (compare Fig. 1B to Fig. 4D, right). We speculated that this failure to recover correlations in the model network, despite the recovery of firing rates, could be the result of perturbing the structured connectivity between excitatory neurons within assemblies generated through training (Fig. 3B). Indeed, the average weights between excitatory neurons within an assembly depressed during the period corresponding to late MD (Fig. S2).

To reveal the origin of this depression in the model network, we investigated the specific contribution of Hebbian plasticity and synaptic scaling to the average weight change within assemblies. Despite the overall potentiation within assemblies induced by synaptic scaling during the period corresponding to early MD, continued LTD from Hebbian plasticity dominated over homeostatic plasticity, depressing all weights within assemblies and preventing the recovery of correlations (Fig. S3). In conclusion, this dominance of depression after MD prevented the recovery of structured connectivity, and consequently correlations, in a model with persistent Hebbian LTD despite homeostatic plasticity. This suggests that the relative timing and resulting competition between the two homeostatic mechanisms and ongoing Hebbian plasticity could be important for recovering different aspects of network dynamics.

### The attenuation of Hebbian LTD together with homeostatic mechanisms restores firing rates and correlations during prolonged MD

Previous work involving ocular dominance plasticity has shown that blocking NMDAR, the main receptor involved in Hebbian plasticity, during the homeostasis-dominant phase causes no significant change in response strength (Toyoizumi et al., 2014). This suggests that the effect of Hebbian plasticity during homeostatic recovery of network activity is negligible. Motivated by this, we asked whether the recovery of correlations during the period corresponding to late MD can be rescued by reducing the effect of Hebbian LTD. We proposed that the attenuation of Hebbian plasticity might occur through a metaplastic process where the amplitude of LTD dynamically adapts to the history of neuronal activity (Methods). To adapt the magnitude of LTD, we followed previous theoretical work (Pfister and Gerstner, 2006; Gjorgjieva et al., 2011). Implementing metaplastic LTD, preserved the recovery of average firing rates of both excitatory and inhibitory neurons (Fig. 5A). Similarly, the spiking rasters during the period corresponding to the early MD phase showed asynchronous irregular activity (Fig. 5B, left). In contrast to the model with persistent LTD, however, the metaplastic reduction in LTD enabled the return of structured activity during late MD (Fig. 5C, left). Importantly, the correlation structure in the model during early and late MD homeostatically recovered after its initial dilution (Fig. 5B and Fig. 5C, right).

**Figure 5:**
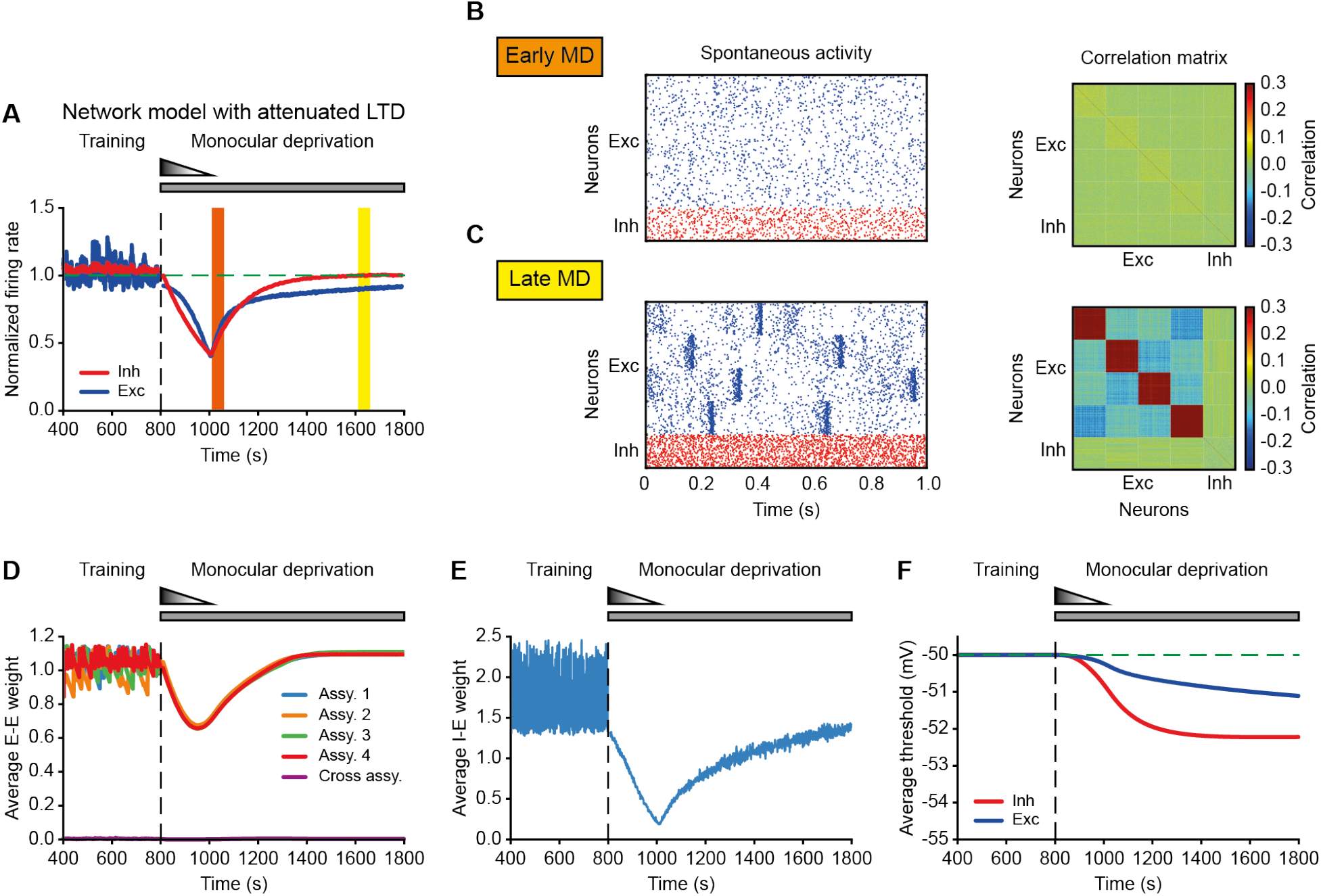
The model with attenuated LTD recovers firing rates and correlations during MD. **A**. The average normalized firing rate of excitatory (blue) and inhibitory (red) neurons. The vertical dashed line indicates the onset of MD. The horizontal dashed line indicates a normalized firing rate of 1.0. **B**. Left: spontaneous activity of excitatory (blue) and inhibitory (red) neurons during early MD. Right: correlation matrix during early MD indicated by the orange region in A. **C**. Same as B but during late MD indicated by the yellow region in A. **D**. Average excitatory-to-excitatory weights for each assembly and across assemblies. **E**. Average inhibitory-to-excitatory weights which target all excitatory neurons independent of assembly membership. **F**. Average firing threshold of excitatory (blue) and inhibitory (red) neurons. The horizontal dashed line indicates the initial firing threshold.

We further investigated what other properties of the network changed as we modeled MD. Along with firing rates and correlations, the average weights within assemblies manifested the same pattern of drop and rebound (Fig. 5D), differently than the weights in the initial model with persistent LTD (Fig. S2). Average inhibitory onto excitatory weights also decreased during early MD in the model (Fig. 5E), suggesting that the network reduced the amount of inhibition to elevate the decreased firing rates of excitatory neurons. During the period corresponding to late MD, overall inhibition increased to balance the gradually recovered excitation, to keep excitatory-inhibitory balance and avoid winner-take-all dynamics where a single strongly-connected assembly dominates the entire network (Litwin-Kumar and Doiron, 2014). Furthermore, the average firing thresholds of excitatory and inhibitory neurons in the model network decreased as we modeled prolonged MD, and reached steady state as the firing rates approached their baseline values (Fig. 5F).

Therefore, metaplastic regulation of LTD together with synaptic scaling and intrinsic plasticity, is sufficient to capture both the recovery of firing rates and correlations during MD. This suggests that homeostatic modifications of overall synaptic weights and intrinsic excitability cooperate with Hebbian LTD to maintain several aspects of network function following input perturbations.

### Individual homeostatic mechanisms have different functionality during MD

To determine the distinct contributions of the different homeostatic mechanisms for the recovery of firing rates and correlations during prolonged MD, we selectively eliminated each mechanism. When deactivating synaptic scaling during the entire period of MD in the model, we found that firing rates still recovered (Fig. 6A), but the correlations did not (Fig. 6C). Since synaptic scaling affects synaptic strengths, we hypothesized that the correlations failed to recover due to the inability of the network to recover its structured connectivity. Indeed, the average weights between excitatory neurons within assemblies remained low in the absence of synaptic scaling (Fig. S4), eliminating structured block activity (Fig. 6B) and preventing the recovery of correlation structure during late MD (Fig. 6C). This suggests that synaptic scaling is indispensable for the recovery of correlations.

**Figure 6:**
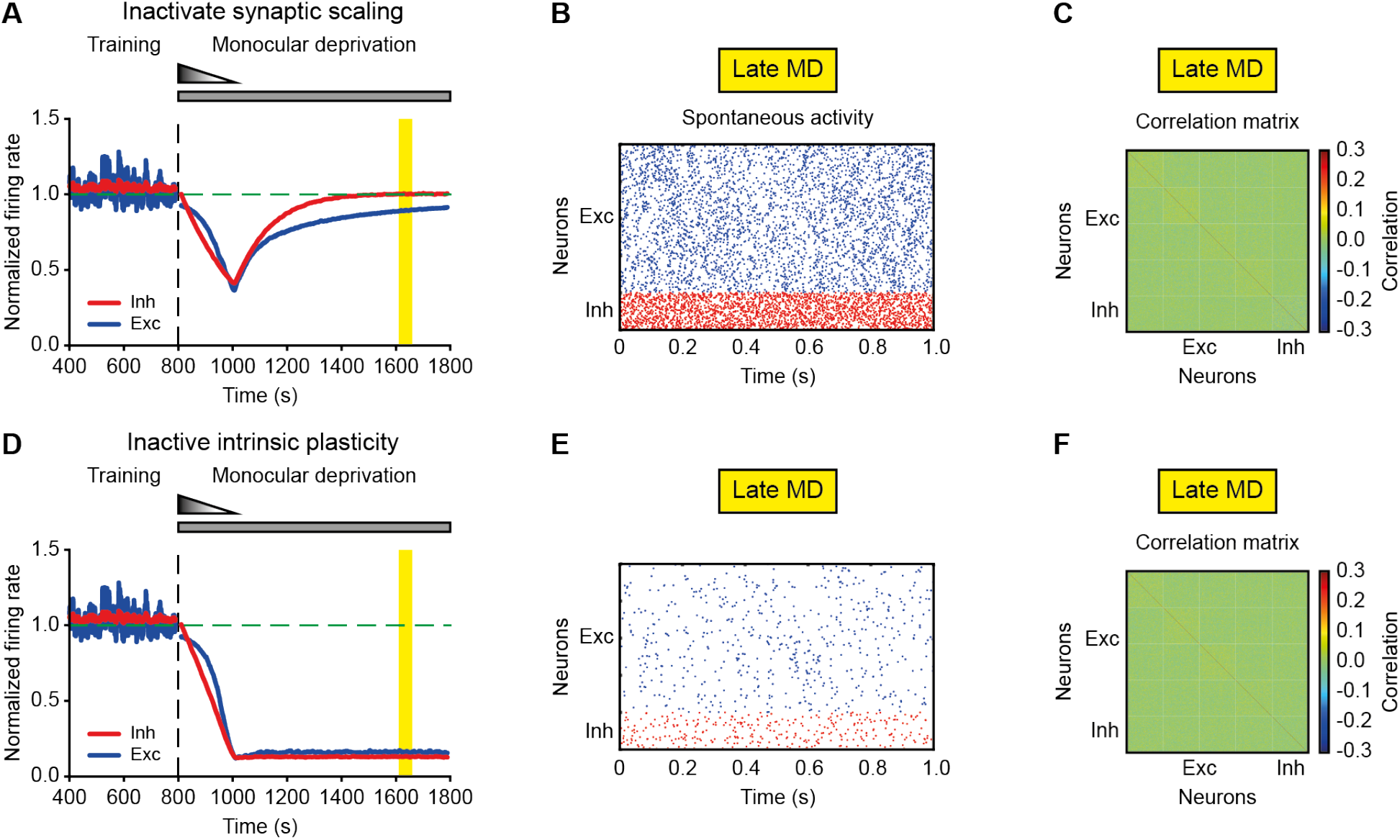
Individual homeostatic mechanisms have different functionality during MD. **A,D.** The average normalized firing rates of excitatory (blue) and inhibitory (red) neurons without synaptic scaling (A) or without intrinsic plasticity (D). The vertical dashed line indicates the onset of MD. The horizontal dashed line indicates a normalized firing rate of 1.0. **B,E.** Spontaneous activity of excitatory (blue) and inhibitory (red) neurons during late MD without synaptic scaling (B) or without intrinsic plasticity (E). **C,F.** Correlation matrix during late MD without synaptic scaling (C) indicated by the yellow period in A, or without intrinsic plasticity (F) indicated by the yellow period in D.

Similarly, without intrinsic plasticity during the entire MD period firing rates in the model did not recover (Fig. 6D). This result was independent of the recovery of correlations. When the overall excitatory drive received by a single neuron within the same assembly was weak, low firing rates were accompanied by a poor degree of synchrony within assemblies (Fig. 6E), resulting in weak correlations (Fig. 6F). Increasing the overall excitation to a neuron, for instance by increasing the connectivity probability within assemblies, generated structured block activity resulting in high correlations within assemblies (not shown), but still failed to recover firing rates to baseline levels, especially for inhibitory neurons.

In conclusion, we demonstrate that two important forms of homeostatic plasticity, synaptic scaling and intrinsic homeostatic plasticity, are able to regulate distinct aspects of network activity. Even though these mechanisms are operating within the context of a specific network architecture implementation here, we propose that they will apply more generally since synaptic and intrinsic plasticity influence different aspects of network function.

## Discussion

A key question in the field of homeostatic plasticity is which aspects of neuronal activity are under homeostatic control. Recent studies have shown that despite a high degree of synaptic plasticity during the critical period (Levelt and Hübener, 2012), firing rates of individual neurons remain remarkably constant during normal development (Hengen et al., 2013), and when perturbed by sensory deprivation rebound back to an individual set point despite continued deprivation (Hengen et al., 2016). Here we used *in vivo* data in rodent visual cortex to investigate whether higher-order cortical network properties are under homeostatic control. We found that – distinct from firing rates – correlations in the control hemispheres increased slightly during early development. In contrast, in deprived hemispheres correlations initially decreased over the first two days and then gradually recovered to pre-deprivation levels, including their structure. Modeling of this process revealed that this restoration of correlation structure could be accomplished through synaptic scaling, while firing rate homeostasis was dependent on intrinsic homeostatic plasticity. Together, these experimental findings provide the first evidence that functional correlation structures are subject to homeostatic regulation.

Recovery of stimulus preference at the single cell level, as well as network correlation structure, has also been reported during repeated episodes of monocular deprivation in the binocular region of visual cortex, each followed by eye reopening (Rose et al., 2016). However, in these ocular dominance plasticity studies, recovery occurring following eye reopening is TrkB-dependent and mediated by Hebbian LTP (Kaneko et al., 2008). This is mechanistically distinct from our work where recovery is governed by homeostatic mechanisms, and where there is no competition between the closed and open eye.

What might be the purpose of the recovered network correlations? Following lid suture to induce MD, the transmitted light through the closed eye lids is relatively weaker compared to the pre-deprivation condition. Therefore, we propose that the network’s homeostatic recovery of correlations might be a way to amplify weak signals, promoting successful signal propagation to other cortical regions (Vogels and Abbott, 2005), which is essential for animals’ perception of the sensory environment (Van Vugt et al., 2018). We predict that the recovery of correlation structure also has important functional implications for information transmission across cortical hierarchies. For instance, neurons in layer 2/3 sum inputs from neurons in layer 4 and are highly influenced by its connectivity and correlation structure. If the recovered network in one layer undergoes a profound remodeling and ends up having a completely different correlation structure, adjustments in successive layers would be needed to keep the cortical network functional.

We cannot conclude from our data whether neurons with higher correlations are more strongly connected. However, as previously shown, functionally correlated neurons are more likely to be connected and if so, more strongly (Ko et al., 2011; Cossell et al., 2015). We therefore assume that correlation strength is indicative of connection strength. In that sense, the identified clusters with strong correlations giving rise to assemblies are consistent with previous experimental work (Ko et al., 2011; Cossell et al., 2015; Barnes et al., 2015). However, this is only the case for excitatory neurons (identified RSUs); since the number of sorted pFS cells was significantly lower than RSUs, we could not investigate their correlation structure.

To dissect the role of various homeostatic mechanisms to restore firing rates and correlations to baseline despite prolonged MD, we built and analyzed a computational model with spiking neurons and biologically realistic plasticity rules. Upon training with correlated input patterns (Ocker and Doiron, 2018), imitating the baseline condition in which animals perceive normal visual inputs, the network exhibited structured spontaneous activity and developed stronger correlations within assemblies. Our model showed that decreasing thalamocortical connection strength (Miska et al., 2018) and decorrelating input patterns during MD, degraded synaptic weights and decreased firing rates and correlations. This was accompanied by a depression in excitatory synaptic weights within assemblies and overall inhibitory synaptic weights in the model. Although experiments have not found significant changes in the strength of recurrent excitation within layer 4 (Maffei et al., 2006), in layer 2/3 there is a generalized depression of excitatory input (Lambo and Turrigiano, 2013); a more systematic analysis that includes measurements within and across assemblies would be necessary to reveal selective depression of some connections.

The modeling results indicated that attenuating the depression effect of Hebbian plasticity was required to maintain clustered network structure during the process of recovery. This suggests that the effect of Hebbian plasticity becomes attenuated during prolonged MD, which then allows homeostatic plasticity to “catch up” and restore network properties. This is consistent with several experimental findings. For example, brief MD leads to occlusion of LTD in layer 4 in primary visual cortex (Crozier et al., 2007; Miska et al., 2018), while homeostatic strengthening of CA1 synapses in hippocampus is accompanied by a reduced ability of synapses to exhibit LTP (Soares et al., 2017). Furthermore, blocking the NMDARs involved in Hebbian plasticity during recovery does not significantly change response strength, suggesting that the total effect of Hebbian plasticity has already attenuated when homeostasis is at its peak (Toyoizumi et al., 2014).

Importantly, in the face of ongoing plasticity, we found that two different forms of homeostatic plasticity can serve distinct functions in recovering network function. First, intrinsic plasticity as a mechanism that affects individual neuron properties, such as the firing threshold, is essential for the rebound of firing rates. Since it does not act directly on the synaptic weights, it has no significant impact on the recovery of correlations. We implemented intrinsic plasticity by simply adjusting the firing threshold which effectively shifts the neuronal input/output function to keep the model sufficiently general. Biophysically, intrinsic plasticity can be implemented by changes in the density and function of voltage-gated channels (LeMasson et al., 1993; Turrigiano et al., 1995; Desai et al., 1999).

Unlike intrinsic plasticity, synaptic scaling regulates synaptic strengths directly and is crucial for the recovery of correlations and network structure in the model. Mechanistically, this regulation is fundamentally distinct from Hebbian plasticity. The regulating process involves an enhanced accumulation of AMPAR in the postsynaptic membrane, which can be mediated by the pro-inflammatory cytokine tumour-necrosis factor-*α* (TNF-*α*) produced by glia (Stellwagen and Malenka, 2006), the immediate-early gene Arc (Shepherd et al., 2006), *β*3 integrins (Cingolani et al., 2008) and other molecules. Crucially, the scaling is bidirectional, global and operates in a multiplicative manner (Turrigiano et al., 1998), although there is some evidence for dendritic-branch specific scaling in some neocortical cell types (Barnes et al., 2017). During recovery, multiplicative scaling potentiates synaptic weights within assemblies more than across assemblies in our model, preserving the relative strength of synaptic inputs and enabling the recovery of correlation structure.

The distinct functional roles fulfilled by synaptic scaling and intrinsic plasticity apply to the context of the present constellation of plasticity rules. We found that synaptic scaling alone is insufficient to recover the firing rates in our model, especially inhibitory firing rates. However, increasing synaptic strengths also boosts neuronal responses, which raises the possibility that synaptic scaling alone might be able to recover firing rates with a different combination of plasticity rules. One straightforward possibility to recover the firing rates of inhibitory neurons is either by increasing the total excitation, for example by upscaling the E-to-I connection, or by decreasing the total inhibition to inhibitory neurons, for example by downscaling the I-to-I connection. Interestingly, synaptic scaling onto inhibitory neurons was recently found to organize model recurrent networks around criticality, independently of firing rates (Ma et al., 2018). This suggests that homeostatic plasticity in excitatory elements might be important for the recovery of firing rates and correlations, while plasticity in inhibitory elements for the recovery of criticality. We highlight that including spiking neurons in our model and training the baseline network with correlated inputs enabled us to study the emergence, dilution and recovery of correlation structure during prolonged MD, which is not possible in unstructured randomly connected networks despite the recovery of firing rates. Furthermore, our implementation of Hebbian and homeostatic plasticity with appropriate biologically motivated timescales suggests a non-trivial cooperation between Hebbian and homeostatic plasticity with the first being attenuated while the latter is in full operation.

In conclusion, our analysis reveals an important, previously unidentified network feature that is homeostatically regulated during perturbation of normal circuit dynamics in the visual cortex. The finding that not only the average correlations, but also the correlation structure, recover has interesting implications for the recovery of computations in these circuits that might be encoded in non-random connectivity patterns. Moreover, our network model with spiking neurons and experimentally characterized homeostatic mechanisms allowed us to dissect the role of each on different aspects of network dynamics, suggesting that different homeostatic mechanisms serve unique, rather than redundant, functions.

## Acknowledgments

We thank all members of the Gjorgjieva lab for comments and discussions. This work was supported by the Max Planck Society (YW, JG), the European Research Council (StG 804824 to JG), RO1 EY025613 and R35 NS111562 (GGT), and R00 NS089800 (KH).

## Author contributions

YW and JG designed the research and developed the model. KH and GGT provided experimental data and contributed to the interpretation of the data analysis. YW analyzed the data with input from all authors. YW performed and analyzed the simulations together with JG. YW and JG prepared the figures and wrote the manuscript with input from GGT.

## Methods

### Firing rates

To obtain the normalized firing rate evolution for different animals, the firing rates of each animal were normalized to the average firing rate at P26 during the light period. Note that here the analysis of firing rates was restricted to MD5 because for the higher order network feature analysis (the pairwise correlations) the number of available, continuously recorded cells beyond this period was insufficient. Therefore, although the firing rates still seem to be above baseline at MD5 – a trend identical to that reported in the previous study (Hengen et al., 2016) – they eventually return to baseline by MD6 (Hengen et al., 2016).

### Pairwise correlations

Each spike train was binned into spike counts of bin size 100 ms, generating a vector of spike counts for each cell. The spike count correlation coefficient *ρ* for a pair of neurons was computed in 30-minute episodes using a sliding window of 5 minutes. We averaged these values for each pair every single half (12-hour) day, thus computing the correlation coefficient for light and dark conditions separately:

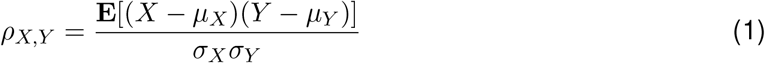

where *X* and *Y* represent the spike count vectors of two cells, respectively; *µ_X_* and *µ_Y_* are the means of *X* and *Y*; *σ_X_* and *σ_Y_* denote the standard deviations of *X* and *Y*; **E** is the expectation. This produced the matrices of pairwise spike count correlations on different half days. Just like the firing rates, to generate the normalized correlation curve across animals, the correlations of each animal were normalized to the average correlation at P26 during the light period.

The correlation matrices in Figure 2A,B were clustered using hierarchical clustering during baseline and the same neuron order was preserved at later time points.

### Similarity quantification of correlation matrices

Shuffled matrix ***A***′ was generated by redistributing the off-diagonal entries of the original matrix ***A*** while keeping the matrix ***A***′ symmetric. The similarity was quantified by computing the absolute difference between the shuffled matrix ***A***′ and the correlation matrix at baseline ***B***:

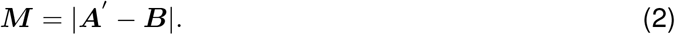

The elements of the upper triangular part of ***M*** were used to form a vector of *L*_1_ distance. Vectors from different animals were then concatenated into a single vector. During shuffling, only the elements corresponding to a given animal were shuffled, i.e. animal identity was preserved.

### Neuron and network model

Single neurons were modeled as leaky integrate-and-fire with membrane potential of neuron *i*, *U*_*i*_, given by (Zenke et al., 2013):

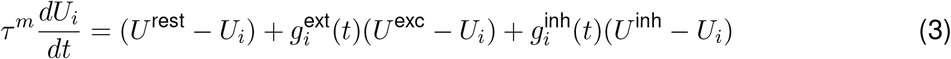

where *τ*^*m*^ is the membrane time constant and *U*^rest^ is the resting potential. The neuron elicited a spike when its membrane potential reached the spiking threshold *U* ^thr^. After a spike, the membrane potential was reset to *U* ^rest^. The neuron also had a refractory period *τ* ^ref^ after a spike. Inhibitory neurons also followed the same integrate-and-fire formalism, but with a shorter membrane time constant. The values of all neuron model-related parameters are listed in Table 1.

The network model consisted of 800 excitatory and 200 inhibitory leaky integrate-and-fire neurons, which were randomly connected with probability of 20%. Excitatory neurons were randomly grouped into four non-overlapping groups. Each excitatory and inhibitory neuron received external excitatory input from 1000 neurons firing with Poisson statistics at average firing rate of 5 Hz, with synaptic strength *w*^ext*→E*^ and *w*^ext*→I*^, respectively.

**Table 1:** Neuron Model Parameters

| Table 1: Neuron Model Parameters |  |  |  |
| --- | --- | --- | --- |
| Symbol | Value | Unit | Description |
| $U^{\text{rest}}$ | -70 | mV | Resting membrane potential |
| $U^{\text{exc}}$ | 0 | mV | Excitatory reversal potential |
| $U^{\text{inh}}$ | -80 | mV | Inhibitory reversal potential |
| $\tau^{\text{ref}}$ | 5 | ms | Duration of refractory period |
| $\tau_{\text{exc}}^m$ | 20 | ms | Membrane time constant of excitatory neurons |
| $\tau_{\text{inh}}^m$ | 10 | ms | Membrane time constant of inhibitory neurons |
| $\tau^{\text{ampa}}$ | 5 | ms | Time constant of AMPA receptor |
| $\tau^{\text{gaba}}$ | 10 | ms | Time constant of GABA receptor |
| $\tau^{\text{nmda}}$ | 100 | ms | Time constant of NMDA receptor |
| $\alpha$ | 0.5 | - | Receptor weighting factor |

Excitatory synapses have a fast AMPA component and a slow NMDA component. Dynamics of excitatory conductances are given by:

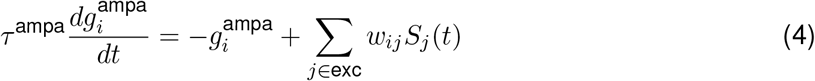

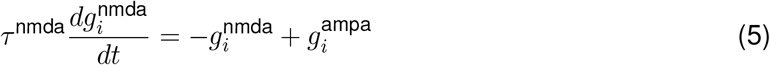

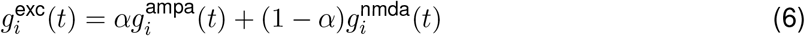

Here *w*_*ij*_ is the synaptic strength from neuron *j* to neuron *i*. If the connection does not exist, *w*_*ij*_was set to 0. *S*_*j*_(*t*) is the spike train of neuron *j*, which is defined as 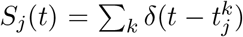, where *δ* is the Dirac delta function 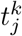 are the spikes times *k* of neuron j. *α* is a weighting parameter.

Dynamics of inhibitory conductances are given by:

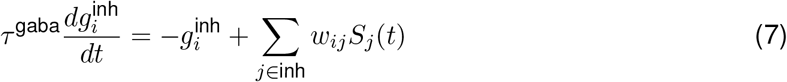

The values of all network-related parameters are listed in Table 2.

**Table 2:** Network Model Parameters

| Symbol | Value | Unit | Description |
| --- | --- | --- | --- |
| $N_E$ | 800 | - | Number of excitatory neurons |
| $N_I$ | 200 | - | Number of inhibitory neurons |
| $p$ | 0.2 | - | Connectivity probability |
| $w^{EE}$ | 0.2 | - | Initial E-to-E connection weight |
| $w^{EI}$ | 2.0 | - | Initial I-to-E connection weight |
| $w^{IE}$ | 0.2 | - | E-to-I connection weight |
| $w^{II}$ | 2.0 | - | I-to-I connection weight |
| $w_{\min}^{EE}$ | 0.0 | - | Minimal E-to-E connection weight |
| $w_{\max}^{EE}$ | 1.2 | - | Maximal E-to-E connection weight |
| $w_{\min}^{EI}$ | 0.0 | - | Minimal I-to-E connection weight |
| $w_{\max}^{EI}$ | 6.0 | - | Maximal I-to-E connection weight |
| $w^{\text{ext} \rightarrow E}$ | 0.78 | - | Initial external-to-E connection weight |
| $w^{\text{ext} \rightarrow I}$ | 0.85 | - | Initial external-to-I connection weight |

### Training procedure

We implemented the network model in three stages: initialization stage, a training stage and an MD stage. All plasticity except for excitatory-to-excitatory plasticity was present in the first 100 seconds of the simulation to initialize the network and obtain network activity before training.

Subsequently, the training process started. During training, correlated stimuli were presented sequentially for 1 second, with 3 second gaps in between stimulus activations. While correlated stimuli were presented to one assembly, the remaining neurons received inputs from 1000 independent neurons firing with Poisson statistics at average firing rate of 5 Hz. The firing rate of the correlated inputs was also 5 Hz. Correlated inputs for the training were generated following previous studies (Brette, 2009; Gjorgjieva et al., 2011). Specifically, we used a copying probability 0.4 from individual uncorrelated Poisson source trains and copying probability 0.6 from a common Poisson source, all with the same firing rates.

The weight matrix obtained after training was used to induce MD in the simulations. MD simulations started with 3 seconds without plasticity when inhibitory STDP was activated, while other plasticity and homeostatic mechanisms were activated at 10 seconds. At the same time, the feedforward connections onto excitatory and inhibitory neurons linearly decreased by 8% and 15% from 10 seconds to 210 seconds. From 210 seconds onwards, feedforward connections were kept fixed.

### Plasticity

To form the clustered correlation structure as observed experimentally, we followed previous modeling studies (Litwin-Kumar and Doiron, 2014) and modeled the plasticity of excitatory-to-excitatory synapses using triplet STDP (Pfister and Gerstner, 2006), of inhibitory-to-excitatory synapses using inhibitory STDP (Vogels et al., 2011) and also heterosynaptic plasticity operating on excitatoryto-excitatory synapses. We used the triplet STDP rule which describes synaptic plasticity based on triplets of spikes and captures experiments where the rate of pre- and postsynaptic neurons varies (Sjöström et al., 2001). The triplet STDP rule enables the formation of bi-directional connections, a necessity for the formation of clustered architectures (Pfister and Gerstner, 2006; Gjorgjieva et al., 2011; Ocker and Doiron, 2018).

According to triplet STDP, the synaptic strength from excitatory neuron *j* to excitatory neuron *i* follows

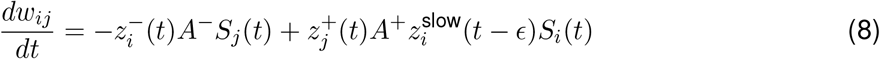

Here, *A^−^* and *A*^+^ are the amplitude of the weight change induced by a post-pre pair or a post-prepost triplet of spikes. *ϵ* is a small positive constant. The synaptic traces 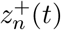, 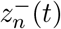 and 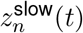 evolved according to

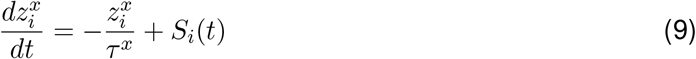

with different time constants *τ*^*x*^.

Unlike excitatory-to-excitatory plasticity, there is significantly less experimental evidence for the form of learning rules operating at other types of synapses. Therefore, inhibitory-to-excitatory plasticity, known as inhibitory STDP, was proposed to counteract the self-feedback loop that potentiates or depresses synaptic strength arising from excitatory-to-excitatory plasticity (Vogels et al., 2011; D’amour and Froemke, 2015). Inhibitory synapses on excitatory neurons are governed by:

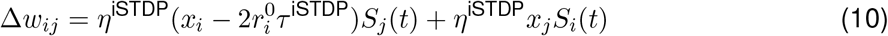

where *x*_*i*_ and *x*_*j*_ are the synaptic traces of the postsynaptic excitatory and presynaptic inhibitory neuron, which are described by 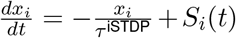 with 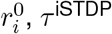 and *η* denoting the target firing rate of neuron *i*, the time constant of the synaptic trace and the learning rate of iSTDP, respectively.

Excitatory-to-inhibitory connections and inhibitory-to-inhibitory connections were non-plastic to reduce the parameters in the model. All plastic weights were subject to upper bounds.

### Heterosynaptic plasticity

We also modeled a second form of normalization in the form of *heterosynaptic plasticity*, which ensures that the sum of all incoming excitatory synaptic weights at each postsynaptic excitatory neuron is kept below a target (Fiete et al., 2010). This form of normalization has been found to be essential in maintaining clustered structures upon their formation (Litwin-Kumar and Doiron, 2014).

The evolution of synaptic strength from excitatory neuron *j* to excitatory neuron *i* via heterosynaptic plasticity follows:

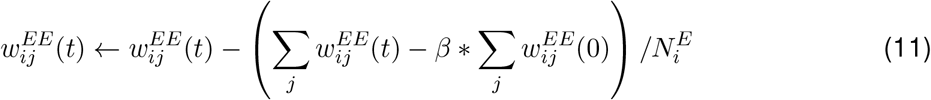

where 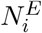 is the number of nonzero elements. As heterosynaptic plasticity also imposed a constraint on the excitatory-to-excitatory synaptic weight, *β* was set to 1.08 so that *w*_*ij*_ becomes approximately 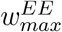. Heterosynaptic plasticity was implemented every 1 s, and only acting when the 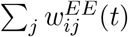 was larger than 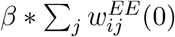.

### Metaplasticity

The amplitude of LTD for neuron i, 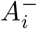 follows

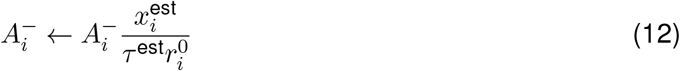

Here, 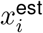 denotes the firing rate estimator defined as 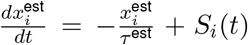 with *τ* ^est^ being the integration time constant of 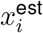. If the firing rate of a neuron was close to its target, 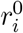, then 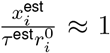. Metaplasticity was implemented every 30 s. Furthermore, 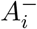 was bounded below by 15% of its initial value to reduce the effect of Hebbian plasticity during the homeostatic recovery phase as shown previously (Toyoizumi et al., 2014).

### Homeostatic mechanisms: synaptic scaling and intrinsic plasticity

The evolution of synapse strength from excitatory neuron *j* to excitatory neuron *i* via synaptic scaling is given by:

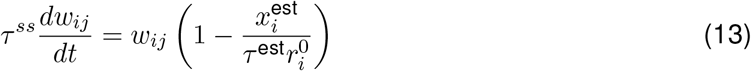

where *τ*^*ss*^ represents the time constant of synaptic scaling.

The firing threshold of neuron *i* regulated by intrinsic plasticity is given by:

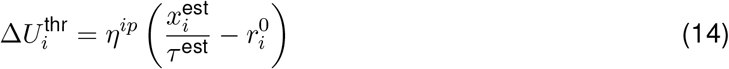

where *η*^*ip*^ is the learning rate of intrinsic plasticity. Initial firing threshold was set to −50 mV. The values of all plasticity-related parameters are listed in Table 3.

**Table 3:** Plasticity Model Parameters

| Symbol | Value | Unit | Description |
| --- | --- | --- | --- |
| $r_0^E$ | 5 | Hz | Target firing rate of excitatory neurons |
| $r_0^I$ | 13 | Hz | Target firing rate of inhibitory neurons |
| $\tau^+$ | 16.8 | ms | Time constant of presynaptic detector |
| $\tau^-$ | 33.7 | ms | Time constant of faster postsynaptic detector |
| $\tau^{\text{slow}}$ | 114 | ms | Time constant of slower postsynaptic detector |
| $A^-$ | 0.0071 | - | Amplitude of LTD |
| $A^+$ | 0.0065 | - | Amplitude of LTP |
| $\tau^{\text{iSTDP}}$ | 0.02 | s | Time constant of synaptic trace for inhibitory STDP |
| $\eta^{\text{iSTDP}}$ | 1 | - | Learning rate of inhibitory STDP |
| $\tau^{\text{est}}$ | 20 | s | Time constant of firing rate estimator |
| $\tau^{\text{ss}}$ | 200 | s | Time constant of synaptic scaling |
| $\eta^{\text{ip}}$ | 0.00025 | - | Learning rate of intrinsic plasticity |

### Simulations

Data analysis and numerical simulations were performed in Python and Julia. All differential equations were implemented by Euler integration with a time step of 0.1 ms.

---

**Figure S1:**
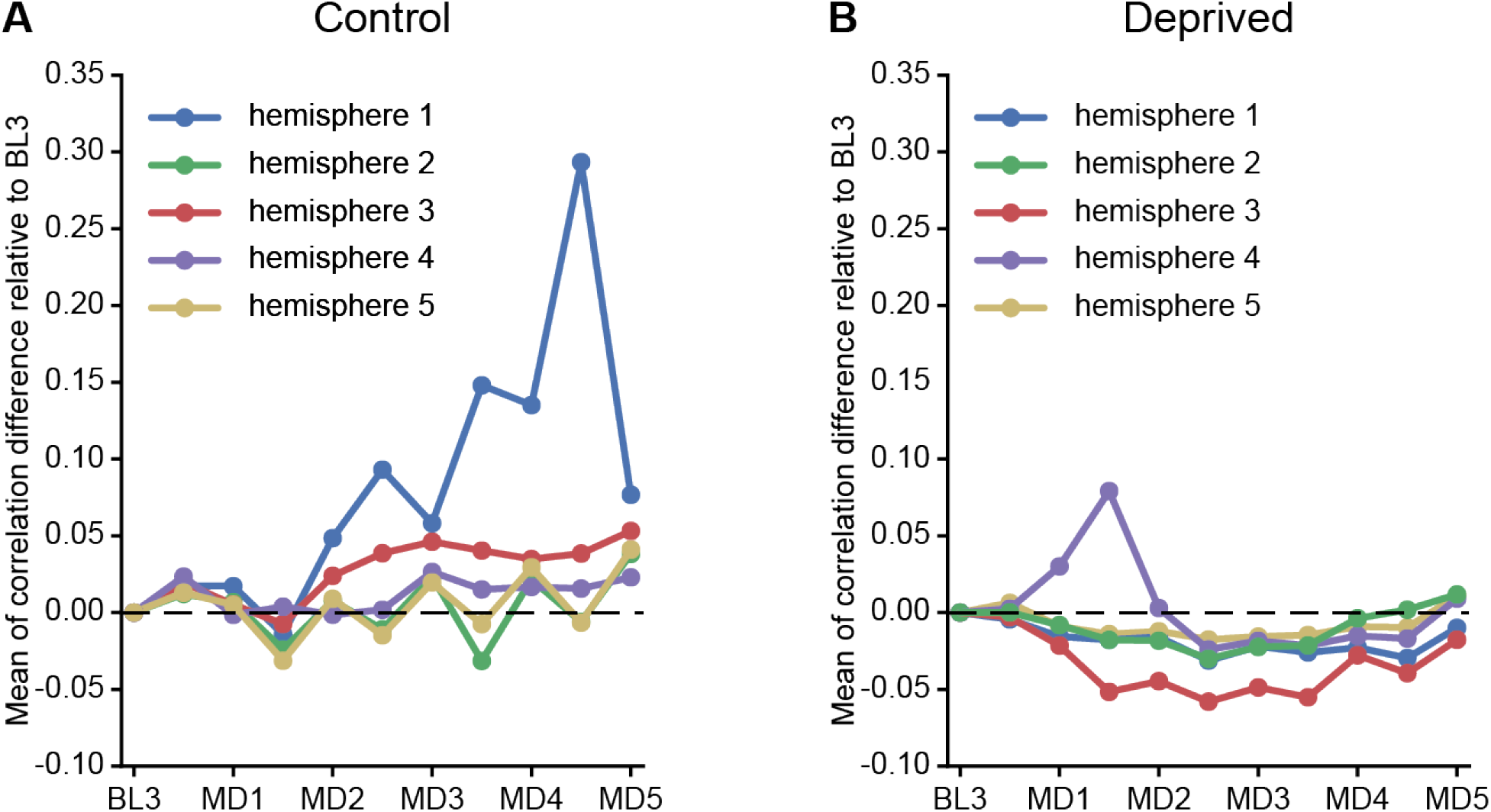
Change in correlations of individual hemispheres. **A**. Correlation differences relative to baseline of five control hemispheres. **B**. Same as A but for five deprived hemispheres. Note that for deprived hemispheres 2 and 4, correlations at MD3 are used for the slope analysis (see Fig. 1E, bottom).

**Figure S2:**
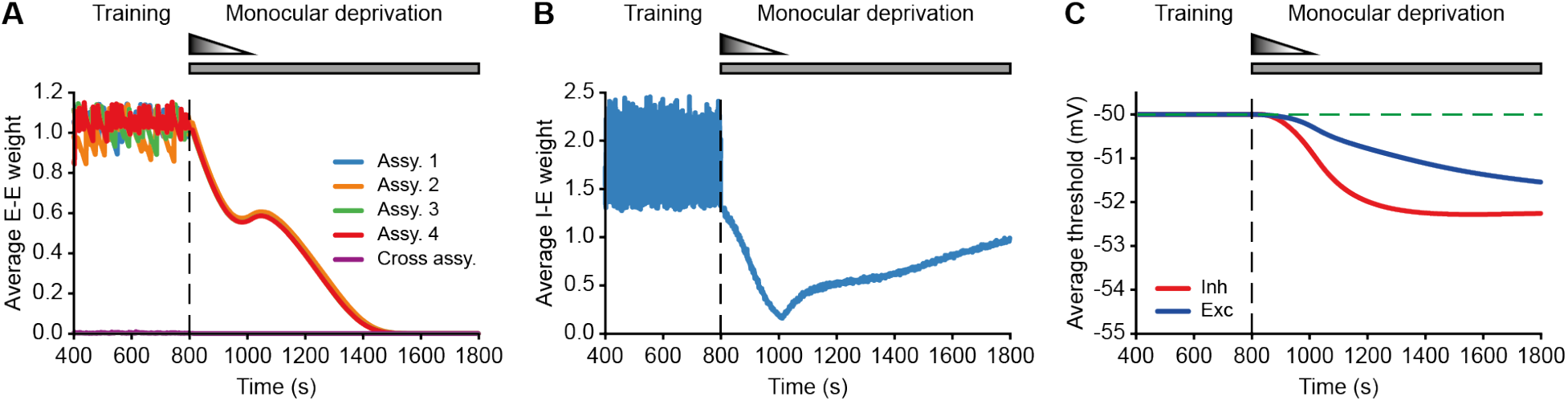
Weights and neuronal threshold in the model with persistent Hebbian LTD and homeostatic plasticity. **A**. Average excitatory-to-excitatory weights for each assembly and across assemblies are depressed by dominant Hebbian synaptic depression. The vertical dashed line indicates the onset of MD. **B**. Average inhibitory-to-excitatory weights which target all excitatory neurons independent of assembly membership. **C**. Average firing threshold of excitatory (blue) and inhibitory (red) neurons. The horizontal dashed line indicates the initial firing threshold.

**Figure S3:**
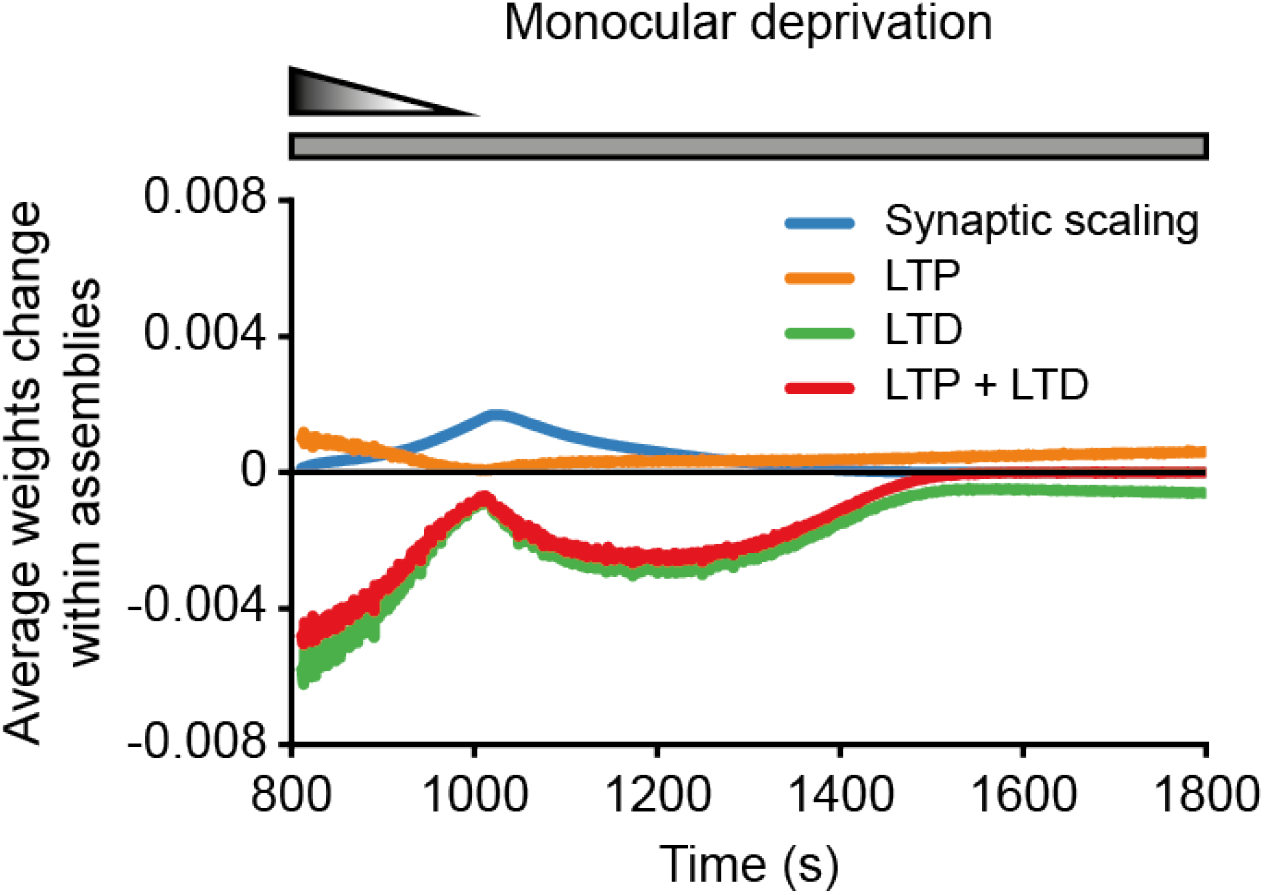
Average weight change within assemblies. Average weight change within assemblies induced by the different synaptic plasticity and homeostatic mechanisms.

**Figure S4:**
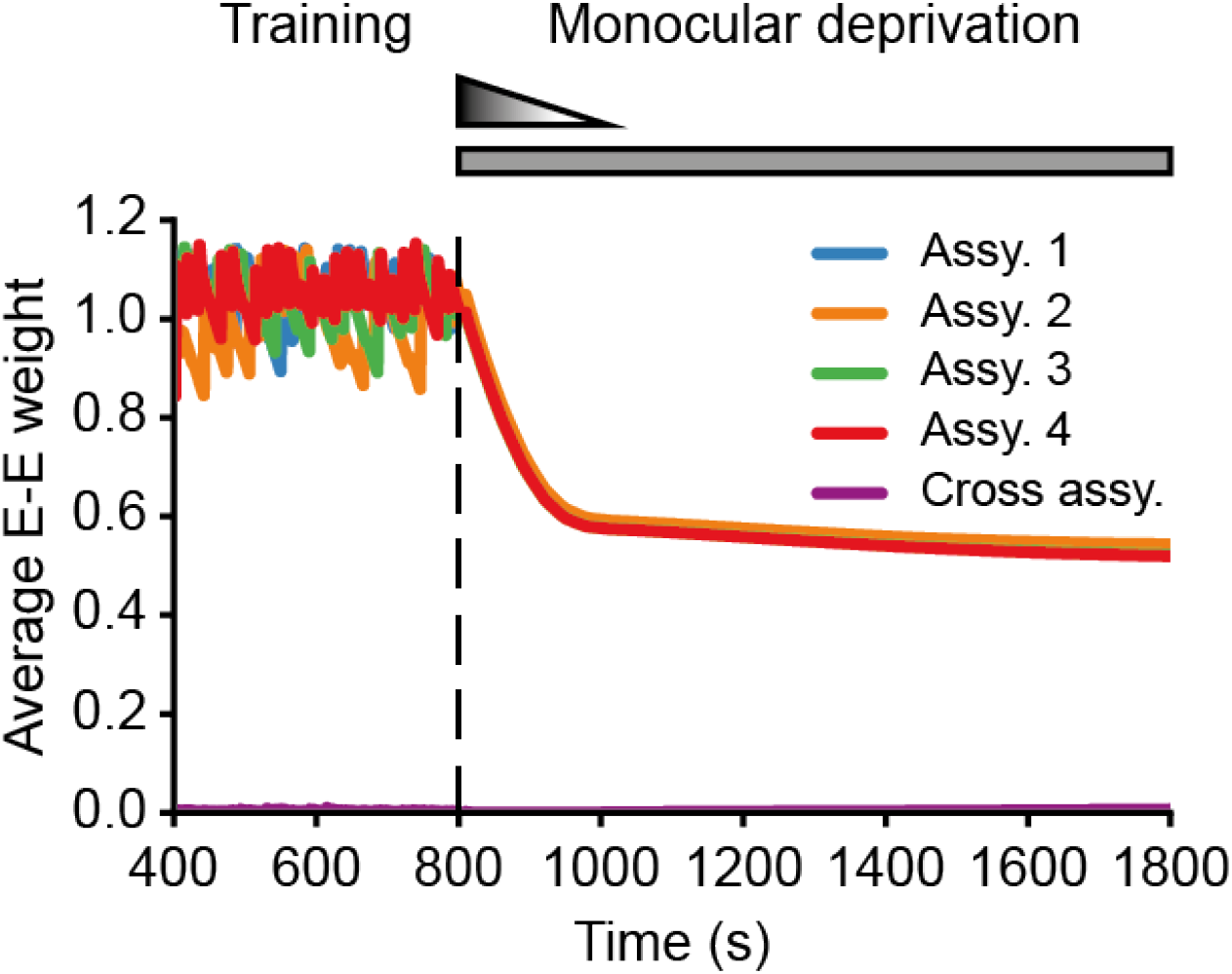
Average E-to-E weight without synaptic scaling. Average excitatory-to-excitatory weights for each assembly and across assemblies without synaptic scaling. The vertical dashed line indicates the onset of MD.

